# hybpiper-rbgv and yang-and-smith-rbgv: Containerization and additional options for assembly and paralog detection in target enrichment data

**DOI:** 10.1101/2021.11.08.467817

**Authors:** Chris Jackson, Todd McLay, Alexander N. Schmidt-Lebuhn

## Abstract

**PREMISE:** The HybPiper pipeline has become one of the most widely used tools for the assembly of target enrichment (sequence capture) data for phylogenomic analysis. Between the production of locus sequences and phylogenetic analysis, the identification of paralogs is a critical step ensuring accurate inference of evolutionary relationships. Algorithmic approaches using gene tree topologies for the inference of ortholog groups are computationally efficient and broadly applicable to non-model organisms, especially in the absence of a known species tree. Unfortunately, software compatibility issues, unfamiliarity with relevant programming languages, and the complexity involved in running numerous subsequent analysis steps continue to limit the broad uptake of these approaches and constrain their application in practice.

**METHODS AND RESULTS:** We updated the scripts constituting HybPiper and a pipeline for the inference of ortholog groups (“Yang and Smith”) to provide novel options for the treatment of supercontigs, remove bugs, and seamlessly use the outputs of the former as inputs for the latter. The pipelines were containerised using Singularity and implemented via two Nextflow pipelines for easier deployment and to vastly reduce the number of commands required for their use. We tested the pipelines with several datasets, one of which is presented for demonstration.

**CONCLUSIONS:** hybpiper-rbgv and yang-and-smith-rbgv provide easy installation, user-friendly experience, and robust results to the phylogenetic community. They are presently used as the analysis pipeline of the Australian Angiosperm Tree of Life project. The pipelines are available at https://github.com/chrisjackson-pellicle.

---

Target enrichment (or sequence capture) is a widely used method for generating high-throughput, multi-locus sequence data for phylogenomic analysis, and it is of greater utility at deeper phylogenetic levels than most other marker systems (McCormack et al., 2013). The approach fragments genomic DNA and then enriches the desired target loci, usually hundreds of genome/gene regions, with RNA baits while removing fragments representing the non-target regions. Bait design consequently requires knowledge of the sequence of the target regions in at least some species of a study group, or closely related species. In recent years an increasing number of bait sets has been designed to enrich either protein coding genes or highly conserved sites flanked by more variable regions (Bejerano et al., 2004; Lemmon et al., 2012) for a variety of major taxonomic groups. In plants, bait sets have been published for flagellate plants (GOFLAG) (Breinholt et al., 2020), flowering plants (PAFTOL / Angiosperms353) (Johnson et al., 2019), Asteraceae (Mandel et al., 2014), mosses (Liu et al., 2019), and ferns (Wolf et al., 2018), among other groups.

Since its publication, the bioinformatics software HybPiper (Johnson et al., 2016) has become one of the most widely used tools for the assembly of target enrichment data (102 citations Web of Science, 166 Google Scholar, accessed 6 June 2021), partly because of its flexibility. It provides options for the assembly of exon or intron sequences, to retrieve a single sequence per sample based on read coverage and contig length, or to collect all potential paralogs for subsequent analysis with other tools using different criteria. A recent adaption of HybPiper developed for the Plant And Fungal Tree Of Life project (Baker et al., 2021), paftools (https://github.com/RBGKew/KewTreeOfLife), does not provide the latter functionality.

The correct inference of ortholog groups is critical in groups showing frequent gene or genome duplication such as many families of land plants, where polyploidy is prevalent. Phylogenetic analysis of paralogous gene copies can produce incorrect topologies, as the evolutionary history of gene families interferes with the evolutionary history of species lineages (Maddison, 1997). Some methods for the inference of ortholog groups require the use of reference genomes (Dessimoz et al., 2012), which remain unavailable in many groups of organisms. Others rely on *a priori* knowledge of ‘undisputed species trees’ (Altenhoff et al., 2016), which creates a conundrum for phylogeneticists, to whom the inference of the species tree is the purpose of the entire exercise. Algorithmic approaches using gene tree topologies to infer ortholog groups, on the other hand, are computationally efficient and have the advantage of broad applicability even in the absence of this kind of data.

A collection of Python scripts published by Yang and Smith (2014) (subsequently Y&S) and recently adapted by Morales-Briones et al. (2020) provides four such algorithms and has become a widely used tool for ortholog inference (107 citations Web of Science, 165 Google Scholar, accessed 6 June 2021). Unfortunately, as originally published, it could not be used on the outputs of HybPiper without reformatting of sequence names and changes to several scripts.

At a practical level, both HybPiper and the Y&S pipeline require the installation of a variety of dependencies on the users’ local system, and the user may be faced with software compatibility issues, creating challenges for the wider adoption of these methods. Moreover, running HybPiper involves five to eight individual terminal commands, and Y&S involves seven to ten (Table 1), depending on the desired results and discounting additional scripts required to pipe HybPiper outputs into Y&S.

**TABLE 1.**
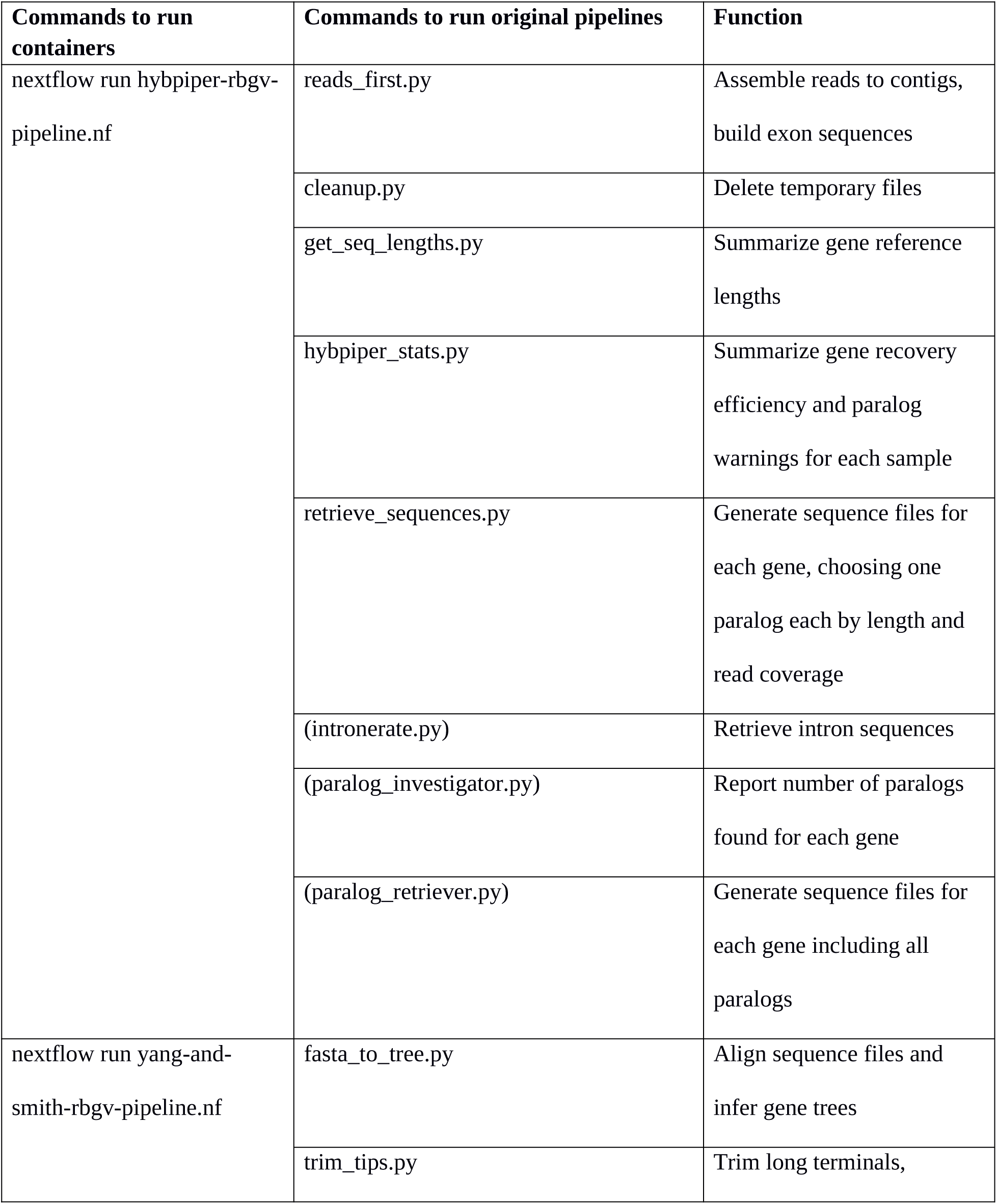
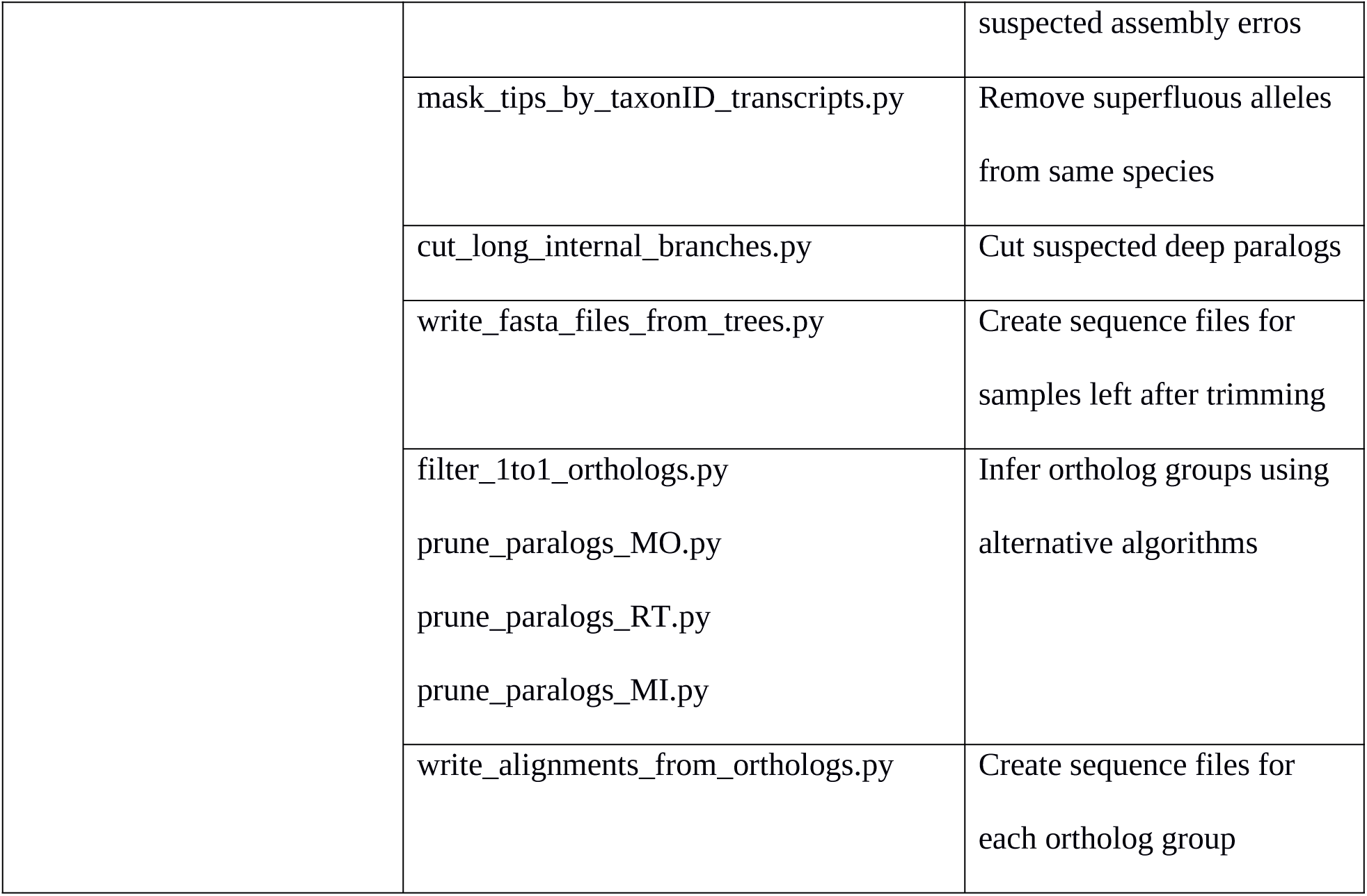
Comparison of command line entries required to run containerized hybpiper-rbgv and yang-and-smith-rbgv against the original implementations of the pipelines, excluding command line arguments. Optional steps are bracketed. Note that additional steps were required to make HybPiper outputs directly usable in the Yang and Smith (2014) pipeline.

To address potential software installation and compatibility issues, we present a Singularity container with all scripts and dependencies required by HybPiper and Y&S pre-installed in a portable software ‘toolbox’. To simplify running HybPiper or Yang and Smith’s (2014) scripts using this container, we provide Nextflow scripts (hybpiper-rbgv and yang-and-smith-rbgv) that allow each improved pipeline to be executed with a single command.

To run hybpiper-rbgv the only inputs required are a folder containing raw reads and a target file in fasta format for the reads to be assembled against. It runs all steps comprising the original HybPiper pipeline, including intronerate and paralog retrieval (https://github.com/mossmatters/HybPiper/wiki/Introns; https://github.com/mossmatters/HybPiper/wiki/Paralogs). One of the outputs of HybPiper are sequence files including all putative paralogs, and these are used as input to the yang-and-smith-rbgv script, together with either a file of outgroup sequences or a list of designated outgroup samples that are already in the HybPiper outputs. The latter outgroup information is required for two of the Y&S ortholog inference algorithms. Additionally, bugs were fixed, and the modified HybPiper code produces more accurate assemblies and flags final locus assemblies that may be built by concatenating SPAdes contigs assembled from different paralogs.

## METHODS AND RESULTS

### hybpiper-rbgv

In the hybpiper-rbgv implementation (Fig. 1), several new features have been added to HybPiper as follows. For each sample, multiple read files (e.g. from different Illumina sequencing machine lanes) can be automatically combined prior to analyses. Input files can now be provided in compressed .gz format. If read quality filtering has not yet been performed, hybpiper-rbgv can optionally run Trimmomatic before assembly. If BLASTx is used for read mapping and the input target file provided contains nucleotide sequences, it is automatically converted to amino acids before prior to BLASTX mapping. If desired, the user can merge forwards and reverse reads prior to assembly using SPAdes.

**FIGURE 1.**
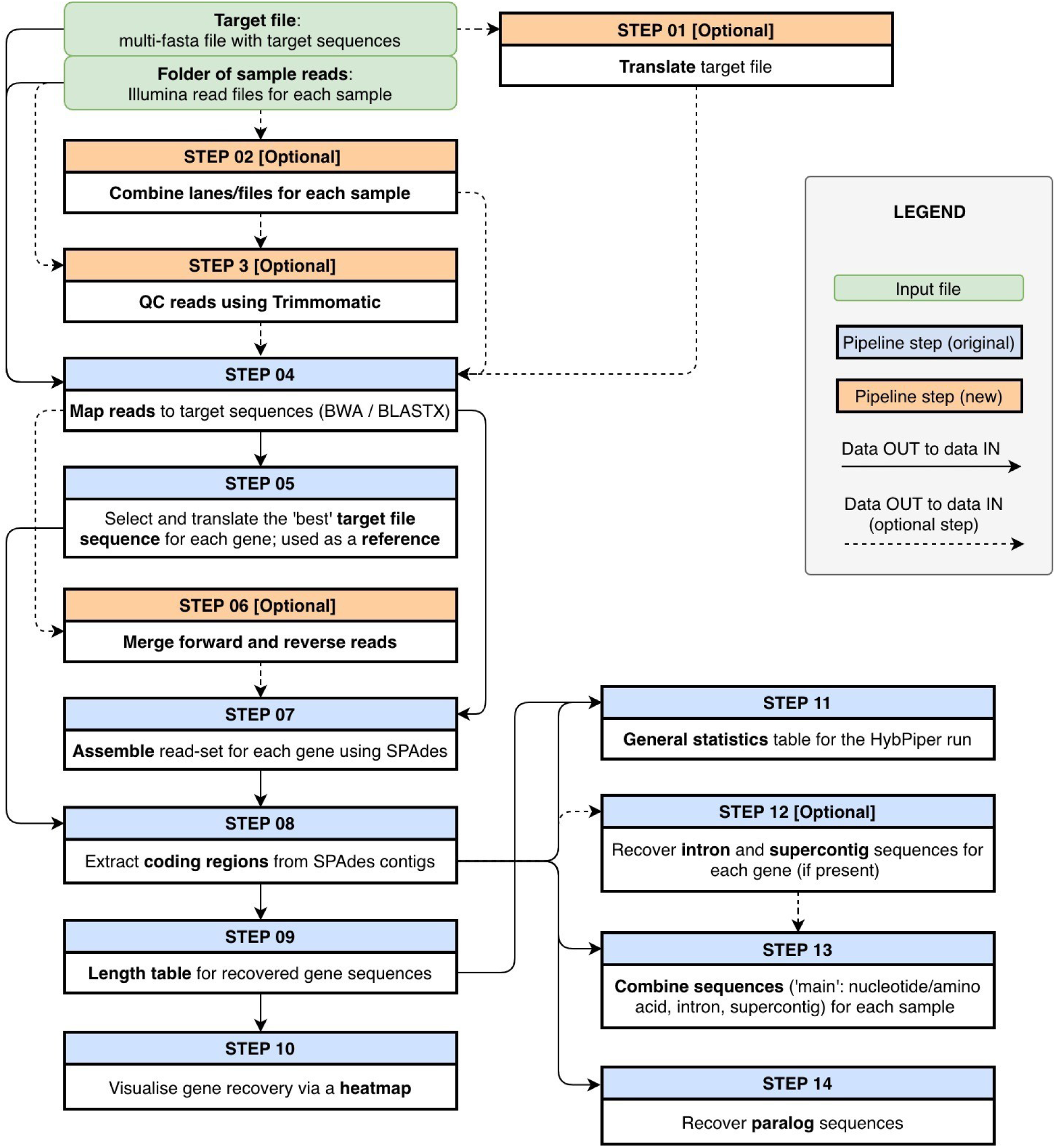
Flowchart summarizing the hybpiper-rbgv pipeline for assembly of sequence capture or target enrichment data.

By default, HybPiper attempts to unite several contigs that individually cover only part of a gene target into a ‘supercontig’. During development we observed that under some circumstances, this approach risks the creation of chimeric supercontigs from different paralogs. Further, supercontig creation can lead to the erroneous duplication of sequence areas at any sites of contig overlap. This latter issue has been fixed in hybpiper-rbgv. To address the former issue, hybpiper-rbgv creates two output folders, one with all supercontigs and one with suspected chimeras (assessed using read-mapping to supercontigs and identification of discordant read-pairs) removed. Optionally, the creation of supercontigs can be suppressed entirely.

In addition, minor bugs were fixed as documented in more detail on the project’s Github site - https://github.com/chrisjackson-pellicle/HybPiper-RBGV.

### yang-and-smith-rbgv

Inference of ortholog groups with the Y&S scripts is based on examination of gene tree topologies. As a first step, the yang-and-smith-rgbv pipeline (Fig. 2) aligns paralog sequences for each gene and infers gene trees. Before the inference of ortholog groups, it conducts trimming of gene trees as implemented in the original pipeline (Yang and Smith, 2014). First, the longer branch in very unbalanced sister terminals is removed, under the assumption that it reveals an assembly or alignment error in the corresponding sequence. Second, very closely related terminals (presumptive alleles) from the same sample are reduced to one, as multiple closely related tips would interfere with the identification of paralogs. Third, very long deep branches are pruned. Minimum parameters for pruning at all steps can be adjusted by the user.

**FIGURE 2.**
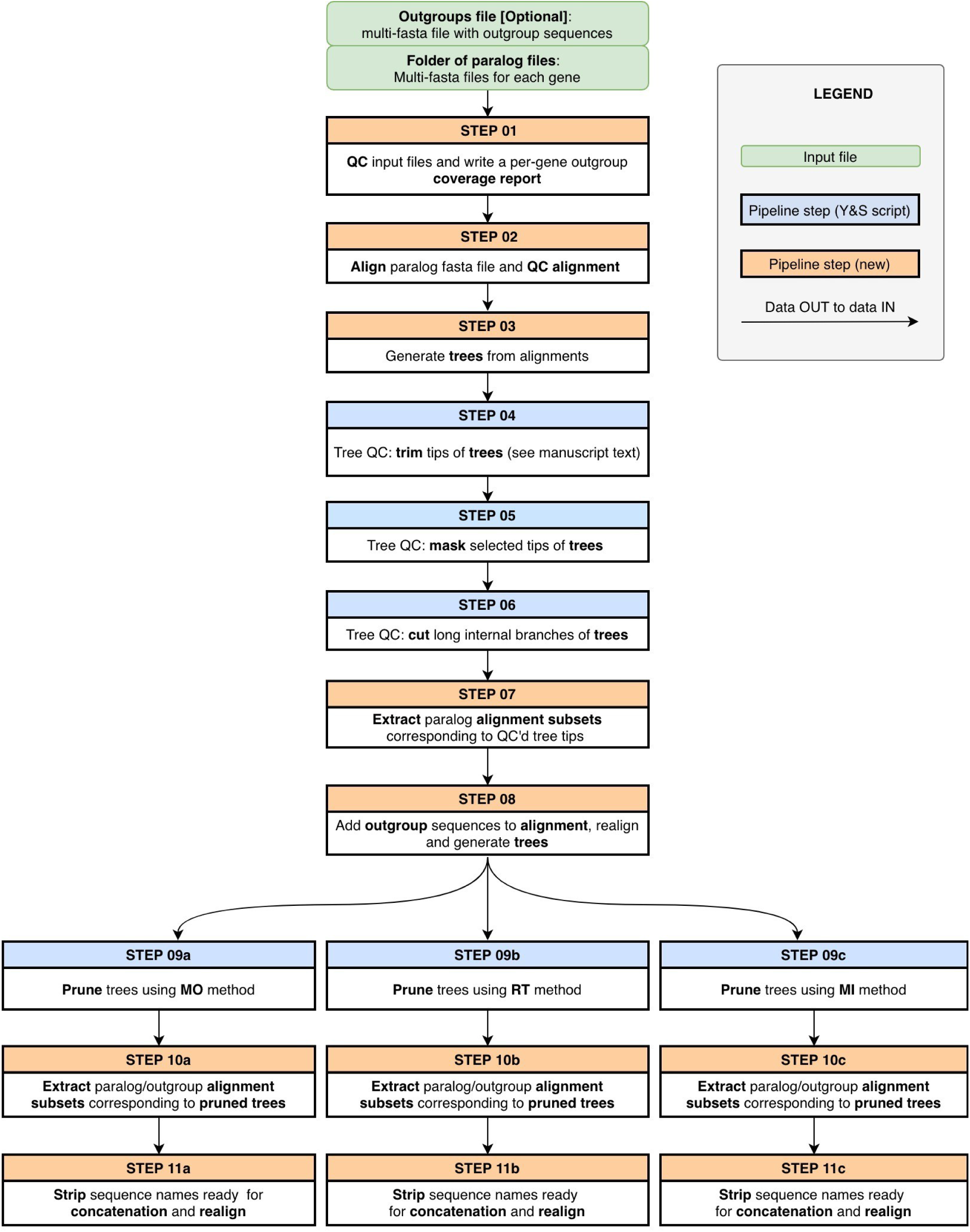
Flowchart summarizing the yang-and-smith-rbgv pipeline, which uses gene tree topology to resolve paralogy, assuming that gene or genome duplication events caused samples to be duplicated in different gene tree clades.

The yang-and-smith-rgbv pipeline implements three of the four algorithms in the collection of Y&S scripts. The Monophyletic Outgroups (MO) algorithm first removes all genes in which the outgroup is non-monophyletic. In the remainder it then iteratively moves upwards from the root, checking at each node if the two daughter clades share samples, and, if so, removes the smaller daughter clade, with the rationale that these nodes represent the location of gene duplication events and that the more informative ortholog group should be kept (Fig. 3a). This approach returns at most the same number of sequence alignments as existed originally.

**FIGURE 3.**
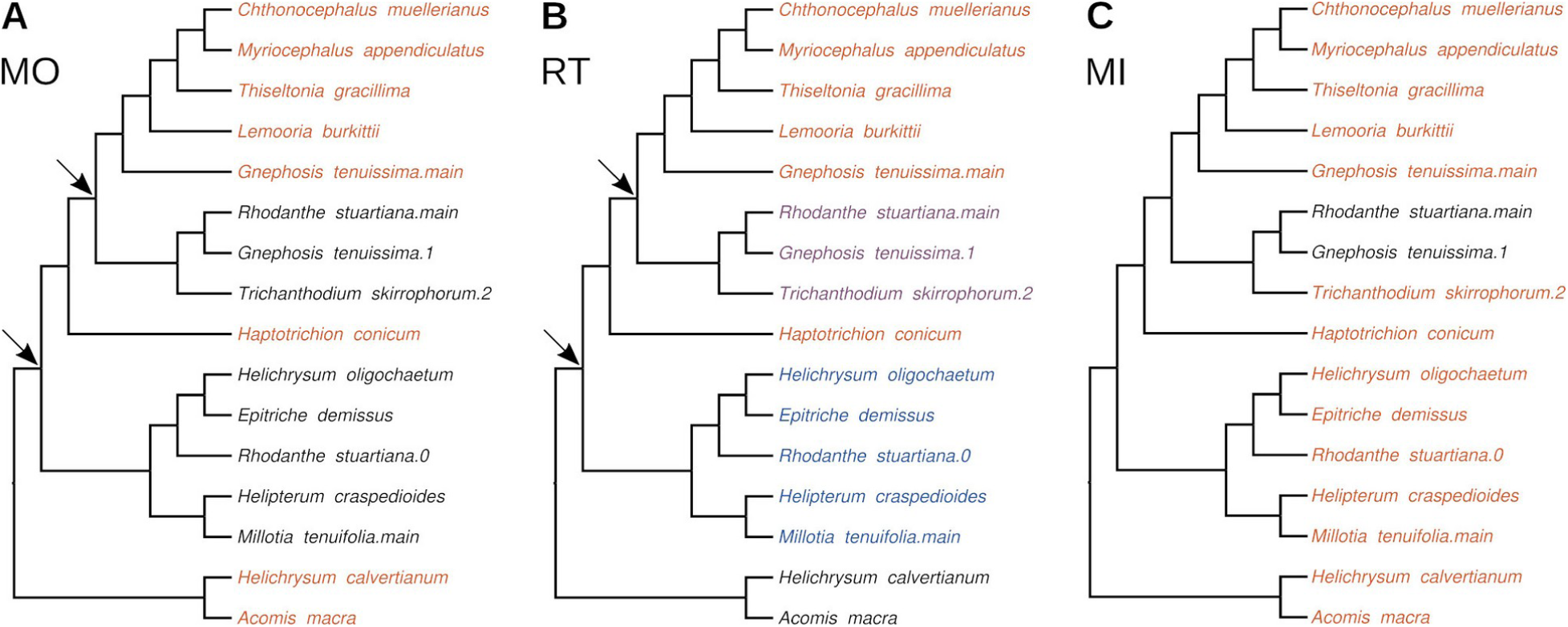
Illustration of algorithms for inference of orthologs using one gene tree as example. (A) Monophyletic Outgroups (MO) moves iteratively through the tree from the root, checks at each node if samples are duplicated between the descendent sister clades, and, if so, removes the smaller descendent sister clade, here retrieving the terminals marked in red. (B) Rooted subTrees (RT) proceeds as MO but separates the smaller descendent sister clades into distinct ortholog groups. In this case, this approach results in the retrieval of three ortholog groups marked in red, blue, and purple. (C) Maximum Inclusion (MI) iteratively retrieves the largest unrooted subtrees without duplicated samples, in this case resulting in a single ortholog group marked in red. The gene tree is presented in cladogram view. Arrows indicate instances of ancestral gene duplication inferred by MO and RT. Name elements after stops are paralog identifiers assigned by HybPiper.

The other two algorithms make use of outgroups supplied as part of the paralog files or in a separate file. Users who need to add outgroups to a dataset from custom baits for which little or no published data are available can mine transcriptome data for sequences matching their HybPiper target file (McLay et al., 2020).

The Rooted subTrees (RT) algorithm first dismantles a gene tree into ingroup clades if the outgroups are non-monophyletic. In each ingroup clade it then iteratively moves upwards from the root, checking at each node if the two daughter clades share samples. If that is the case, it separates the smaller daughter clade out as a new ortholog group under the assumption that a gene duplication occurred at this node (Fig. 3b). Consequently, this approach has the potential to output considerably more sequence files than in the original input, and some ortholog groups may contain very few samples.

The Maximum Inclusion (MI) algorithm iteratively extracts the largest subtrees from an unrooted gene tree that do not contain duplicated samples (Fig. 3c). In contrast to MO and RT, this approach does not rely on a logic that locates putative gene duplication events and may consequently be considered less theoretically defensible than the alternatives.

The final algorithm of Yang and Smith (2014), 1to1, simply removes all genes containing paralogs, retaining only the paralog-free genes. While this not explicitly implemented in yang-and-smith-rbgv, the user can select all files labeled ‘1to1ortho’ from the results of the Maximum Inclusion algorithm to achieve the same outcome.

The yang-and-smith-rbgv pipeline produces gene alignments for each inferred ortholog group under each of the three algorithms. These alignments are ready for phylogenetic analysis either separately or after concatenation. The pipeline uses MAFFT v. 7.471 (Katoh and Standley, 2013) or MUSCLE (Edgar, 2004) for alignment steps and IQ-TREE v. 2.0.3 (Nguyen et al., 2015) for gene tree inference.

### Example dataset

We tested the two pipelines on several datasets predominantly of Asteraceae and Orchidaceae. Most analyses used the Angiosperms353 bait set (Johnson et al., 2016), and one used the compositae1061 bait set (Mandel et al., 2014). A small dataset of twelve ingroup and two outgroup Asteraceae is here used as an example. It is drawn from tribe Gnaphalieae: subtribe Gnaphaliinae: Australasian clade (Schmidt-Lebuhn and Bovill, 2021). The data were produced by the Australian Angiosperm Tree of Life project as part of the Genomics for Australian Plants consortium (https://www.genomicsforaustralianplants.com/). Raw reads were quality filtered and trimmed using Trimmomatic 0.38 (Bolger et al., 2014). Only paired reads were used for subsequent assembly with hybpiper-rbgv (though the input can include single orphan reads from a Trimmomatic run, as well as a new option to include merged reads). The target file for assembly was produced by filtering the angiosperm megatarget file of McLay et al. (2020) for Asteraceae. Ortholog groups were inferred for resulting sequence files including paralogs (‘11_paralogs’ directory) using all algorithms implemented in yang-and-smith-rbgv under default settings. For the MO and RT algorithms, *Acomis macra* F.Muell. and *Helichrysum calvertianum* (F.Muell.) F.Muell. were set as outgroups. They were selected because they belong to the Waitzia clade of Australasian Gnaphalieae (Schmidt-Lebuhn and Bovill, 2021). In each case, we removed genes or ortholog groups with data for less than five samples.

Sequence alignments for each ortholog group were processed to ensure that they were all in the correct frame and concatenated using custom Python scripts. We compared dataset characteristics and phylogenetic results for five different approaches: the results from each algorithm for inference of ortholog groups (MO, RT, MI); only the paralog-free genes; and the direct HybPiper output, which selects a paralog to maximise contig length and read coverage. In each case, we reconstructed a species tree using ASTRAL 5.7.7 (Zhang et al., 2018) after inferring individual gene trees with IQ-TREE 1.6.12 (Nguyen et al., 2015) under the HKY+I+G model, also partitioning by codon position.

### Comparison of ortholog inference approaches

After filtering for read quality, the 14 samples in the example dataset retained 1,007,159 to 40,976,703 reads (median 5,895,305). Of these, between 5.1% and 56.1% were on-target (median 28.2%). hybpiper-rbgv retrieved sequences for between 296 and 348 genes (median 342) per species, of which between 166 and 283 (median 251) were at least 75% of the length of the mean length of all target sequences for a given gene. In total, hybpiper-rbgv produced gene files for 350 of the 353 targeted genes. Between 9 and 29 genes (median 20) generated paralog warnings; HybPiper statistics are available at DOI: 10.25919/q42q-j056.

Dataset sizes are summarised in Table 2. Using the outputs of hybpiper-rbgv directly resulted in 296-345 genes per species (median 340.5), as five genes were excluded for having less than five terminals. The MO algorithm of yang-and-smith-rbgv removed 51 genes for having non-monophyletic outgroups, removed paralogs from 22 genes, and inferred no paralogs in 277 genes, for a total of 299 remaining ortholog groups.

**TABLE 2.** Dataset sizes resulting from different algorithms for the inference of ortholog groups in a test dataset of fourteen Australian Asteraceae. In larger datasets, the use of paralog-free genes only is likely to result in relatively smaller datasets, and that of the Rooted subTrees algorithm in relatively larger ones.

| Algorithm | Ortholog groups per species, min-max (median), after filtering for ≥5 terminals | Characters |  |  |  |
| --- | --- | --- | --- | --- | --- |
|  |  | Total | Parsimony informative | Variable but uninformative | Constant |
| No ortholog inference | 296-345 (340.5) | 273,042 | 34,485 | 49,600 | 188,957 |
| Monophyletic Outgroups | 245-293 (286) | 210,090 | 27,836 | 36,530 | 145,724 |
| Rooted subTrees | 224-253 (235) | 209,613 | 19,933 | 34,319 | 155,361 |
| Maximum Inclusion | 306-352 (348) | 251,499 | 32,095 | 42,815 | 176,589 |
| Only paralog-free genes | 229-273 (268) | 195,822 | 25,883 | 34,239 | 135,700 |

The RT algorithm inferred the existence of 642 ortholog groups but only resulted in 224-253 ortholog groups per species carried over into phylogenetic analysis (median 235), because 335 ortholog groups were excluded for having data for less than five species.

The MI algorithm inferred no paralogs for 277 and separated 139 ortholog groups out of the remaining 73, for a total of 416 resulting ortholog groups. It resulted in 306-352 ortholog groups per species (median 348), with 36 ortholog groups excluded for having less than five terminals.

Using only paralog-free genes resulted in 229-273 genes per species (median 268), with 3 genes excluded for having less than five terminals.

The ASTRAL phylogeny inferred for direct HybPiper outputs without inference of ortholog groups differs from that inferred for all ortholog inference approaches in the relationships of *Chthonocephalus muellerianus* P.S.Short, *Epitriche demissus* (A.Gray) P.S.Short, *Gnephosis tenuissima* Cass., and *Trichanthodium skirrophorum* Sond. & F.Muell. ex Sond., suggesting that the analysis is misled by the presence of unrecognized paralogy (Fig. 4). In addition, the placement of *Millotia tenuifolia* Cass. varies across analyses, with data derived from the MI and RT algorithms favoring one placement, and those from MO and only paralog-free genes another.

**FIGURE 4.**
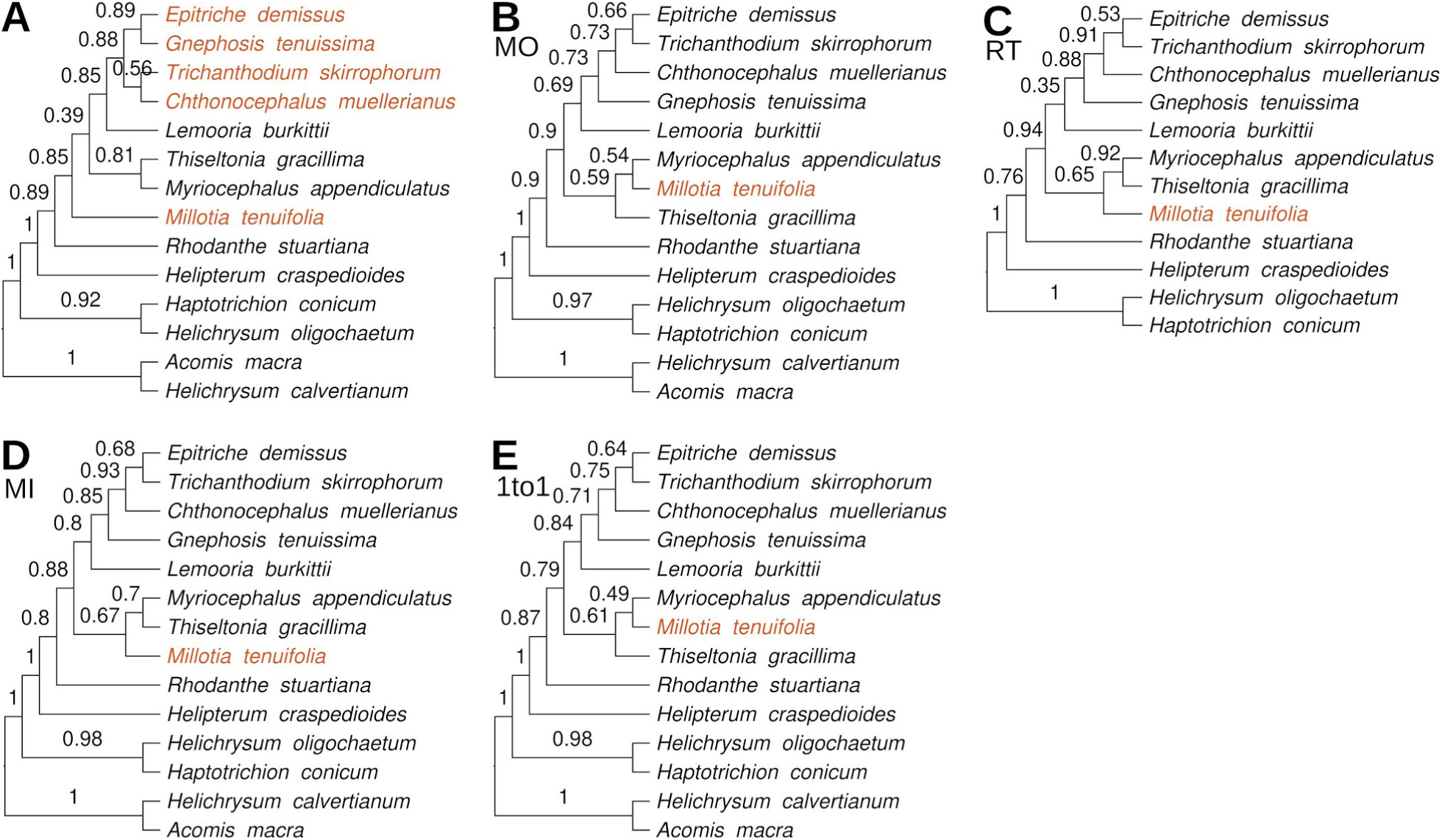
Results of phylogenetic analysis of the example dataset with ASTRAL, using data from orthology inference by (A) hybpiper-rbgv directly, based on length and read coverage, (B) Monophyletic Outgroups, (C) Rooted subTrees, (D) Maximum Inclusion, and (E) exclusion of all genes with paralogs. Outgroup is missing in (C) because the RT algorithm removes it. Numbers above branches indicate clade support from local posterior probability. Red font color marks a species placed in differing positions and a clade whose internal structure differs in (A), whereas the remainder of the topology is constant across analyses.

## CONCLUSIONS

hybpiper-rbgv and yang-and-smith-rbgv are pipelines for the assembly of target enrichment data and the inference of ortholog groups that facilitate installation and simplify use compared to the standalone HybPiper and Yang-and-Smith softwares. They required little to no expertise in scripting and provide several new options, increasing flexibility with regard to input data e.g. by allowing the use of read files from multiple lanes.

By improving the method of joining contigs from the same gene together, hybpiper-rbgv does not produce duplicated sequence regions during the generation of supercontig-derived loci sequences. Additionally, it implements options for the removal of potentially chimeric supercontigs or of all supercontigs, giving the user additional assembly options. yang-and-smith-rbgv implements the same algorithms for ortholog inference as its original version but can use the outputs of hybpiper-rbgv directly and provides greater flexibility for the use of outgroups.

Our testing of the algorithms implemented by Yang and Smith (2014) across different datasets, here exemplified with a set of fourteen Australian Asteraceae, illustrated the benefit of the removal of paralogs, the benefit of including genes exhibiting paralogy, and the relative performance of the topology-based approaches. The phylogeny inferred without formal ortholog resolution deviated from all others, suggesting that its topology is influenced by unrecognised paralogy (Fig. 4a). Removing all genes showing paralogy, however, produced the smallest dataset, albeit with slightly more informative characters than the results of RT (Table 2). This effect would be stronger in larger datasets, as the number of gene files containing at least one paralog increases with the number of species in the analysis.

Similarly, the number of species with paralogs will increase with the number of genes, and vice-versa. As expected, Maximum Inclusion (MI) produced the largest paralog-free dataset, and the resulting phylogeny was not an outlier among those derived from the paralog-free datasets (Fig. 4b). Rooted subTrees (RT) separated out the largest number of ortholog groups but resulted in the smallest dataset after filtering for a minimum number of terminals per ortholog group, an artefact of the small size of the example dataset. In larger test datasets, this approach frequently produced more informative datasets than Monophyletic Outgroups (MO) (Schmidt-Lebuhn, unpubl. data).

Depending on the data, additional processing may be desirable before phylogenetic analysis, e.g. to ensure that all genes are in the correct frame if protein-coding. Nevertheless, hybpiper-rbgv and yang-and-smith-rbgv greatly streamline the assembly of target enrichment data and inference of ortholog groups, making these methods more accessible and easier to use by those working with target capture dataset.

## ACKNOWLEDGMENTS

We are grateful to Matt G. Johnson, Stephen A. Smith, and Ya Yang for discussions on and clarifying the use of their pipelines, and to Theodore Allnut, Jason Bragg, Johan Gustafsson, Cameron Jack, and Lars Nauheimer for helpful conversations and feedback during development. Sequencing was funded by Bioplatforms Australia through the Genomics for Australian Plants (GAP) initiative as part of the phylogenomics project Australian Angiosperm Tree of Life (AAToL). We used the sequencing services provided by the Australian Genome Research Facility (AGRF).

## DATA AVAILABILITY

The hybpiper-rbgv and yang-and-smith-rbgv containers are available at https://github.com/chrisjackson-pellicle. The example dataset, HybPiper statistics, target file, and outgroup file are available at the CSIRO Data Access Portal (DOI:10.25919/q42q-j056). The raw reads of the example dataset are available in the Bioplatforms Data Portal (https://data.bioplatforms.com/) under sample numbers 79649, 79652, 80014, 80042, 80066, 80070, 80071, 80082, 80088, 80089, 80105, 80109, 80123, and 80125.

